# Loss of Caveolin-1 and caveolae leads to increased cardiac cell stiffness and functional decline of the adult zebrafish heart

**DOI:** 10.1101/2020.01.16.909267

**Authors:** Dimitrios Grivas, Álvaro González-Rajal, Carlos Guerrero Rodríguez, Ricardo Garcia, José Luis de la Pompa

## Abstract

Caveolin-1 is the main structural protein of caveolae, small membrane invaginations involved in signal transduction and mechanoprotection. Here, we generated *cav1-KO* zebrafish lacking Cav1 and caveolae, and investigated the impact of this loss on adult heart function and response to cryoinjury. We found that cardiac function was impaired in adult *cav1-KO* fish, which showed a significantly decreased ejection fraction and heart rate. Using atomic force microscopy, we detected an increase in the stiffness of epicardial cells and cortical myocardium lacking Cav1/caveolae. This loss of cardiac elasticity might explain the decreased cardiac contraction and function. Surprisingly, *cav1-KO* mutants were able to regenerate their heart after a cryoinjury but showed a transient decrease in cardiomyocyte proliferation.

## Introduction

Caveolae are small membrane invaginations present in endothelial cells, fibroblasts and less abundantly, in cardiomyocytes (Cohen et al., 2003; Patel et al., 2007; Robenek et al., 2008; Scherer et al., 1997). Caveolin-1 (Cav1) is the main structural protein of caveolae (Fra et al., 1995), as *Cav1* deletion in mice abolishes caveolae formation (Drab et al., 2001). Similarly, the caveolae associated protein Cavin1, is essential for caveolae formation, because its genetic deletion leads to loss of caveolae (Liu et al., 2008). Caveolae participate in multiple cellular processes, including lipid homoeostasis and signal transduction (Cheng et al., 2015; Cohen et al., 2003; Kim et al., 2008). In particular, Cav1 interacts directly with TGFβR1, blocking Smad complex nuclear translocation and, consequently, inhibiting transcriptional activation (Razani et al., 2001). Furthermore, caveolae are involved in mechanoprotection, as they deliver the extra membrane needed for cells to buffer the mechanical forces through rapid disassembly and flattening (Sinha et al., 2011). Physiologically, caveolae protect mouse cardiac endothelial cells from rupture caused by increased cardiac output (Cheng et al., 2015). Likewise, caveolae safeguard zebrafish skeletal muscle cells from rupture after vigorous activity (Lo et al., 2015), and maintain notochord’s integrity (Garcia et al., 2017a; Lim et al., 2017).

Knock-out of *Cav1* in the mouse results in cardiac remodelling. Right ventricle dilatation and left ventricle hypertrophy are among the various cardiac defects associated with loss of caveolae (Cohen et al., 2003; Park et al., 2003; Zhao et al., 2002). Additionally, *Cav1* mutant mice show defective heart function, including decreased systolic and diastolic function (Cohen et al., 2003; Park et al., 2003; Wunderlich et al., 2006; Zhao et al., 2002), which are exacerbated after myocardial infarction (Jasmin et al., 2011; Shivshankar et al., 2014). Cardiac insult in *Cav1* mutant mice also leads to aberrant fibrosis, mediated by increased Smad2/3 phosphorylation and M2 macrophages infiltration (Miyasato et al., 2011; Shivshankar et al., 2014). Resection of the ventricular apex in hearts of *cav1a* mutant zebrafish leads to regeneration arrest 30 days post injury, because of decreased cardiomyocyte proliferation and increased fibrosis in the amputation plain (Cao et al., 2016).

Here, we generated *cav1-KO* zebrafish and investigated the importance of Cav1 and caveolae in the mechanical properties of the cardiac tissue and in regeneration after cryoinjury. We found that while the absence of Cav1 does not affect cardiac regeneration, *cav1-KO* hearts show a transient decrease in cardiomyocyte proliferation during this process. Using atomic force microscopy (AFM)-force spectroscopy measurements (Dufrêne et al., 2017) we detected a significant decrease in cardiac elasticity in *cav1-KO* animals. Accordingly, epicardial cells and cortical myocardium in *cav1-KO* hearts lacking caveolae are stiffer than wild type (WT) counterparts. Furthermore, *cav1-KO* hearts showed a severe ventricular dysfunction, underscoring the role of caveolae in the mechanical properties and homeostasis of the heart.

## Results and Discussion

### Caveolin-1 expression in the intact and regenerating zebrafish heart

We began our analysis by examining Cav1 expression in intact hearts. We used the *Tg(fli1a:GFP)* line (Lawson and Weinstein, 2002), which expresses GFP in the endocardium and endothelium, and stained with antibodies against Cav1 and tropomyosin (Fig. 1A). Robust Cav1 expression was detected in the vasculature (asterisks in Fig. 1B-B΄) and in the endocardium (arrowheads in Fig. 1B-B΄ and Fig. 1C-C΄). Strong expression was also found in the epicardium (arrows in Fig. 1B-B΄), in the bulbus arteriosus and in the valves (Fig. 1A). Additionally, Cav1 expression was detected in the area between the cortical and trabecular myocardium (dashed area in Fig. 1B-B΄ and inset in Fig. 1B΄). We then analysed Cav1 expression in the regenerating zebrafish heart after cryoinjury (Fig. 1D). We used the *Tg(wt1b:GFP)* line that expresses GFP in the epicardium upon injury (González-Rosa et al., 2012). Seven days post cryoinjury (dpci), Cav1 was strongly expressed in epicardial cells (Fig. 1E, brackets) covering the injured site, overlapping with GFP. High expression was also detected in the endocardium within the injured area (Fig. 1E, arrows). To confirm these observations, we utilised the *Tg(fli1a:GFP)* line and found that Cav1 was expressed in GFP^+^ endocardial cells invading the damaged tissue (Fig. 1F-G΄, arrows). We also surveyed the expression of caveolae-related genes during heart regeneration by quantitative (q)PCR (Fig. 1H). *cav1* and *cavin1b* were upregulated after injury, in contrast to *cav2* and *cav3* whose expression remained stable. These results show that Cav1 is expressed in the endocardium, endothelium and epicardium of the intact heart, three cell types that are activated during regeneration (González-Rosa et al., 2012; Kikuchi et al., 2011; Marín-Juez et al., 2016; Münch et al., 2017). Also, upon injury, Cav1 expression is strongly increased in epicardial cells surrounding the injured site, and in the endocardium invading the injured area.

**Figure 1.**
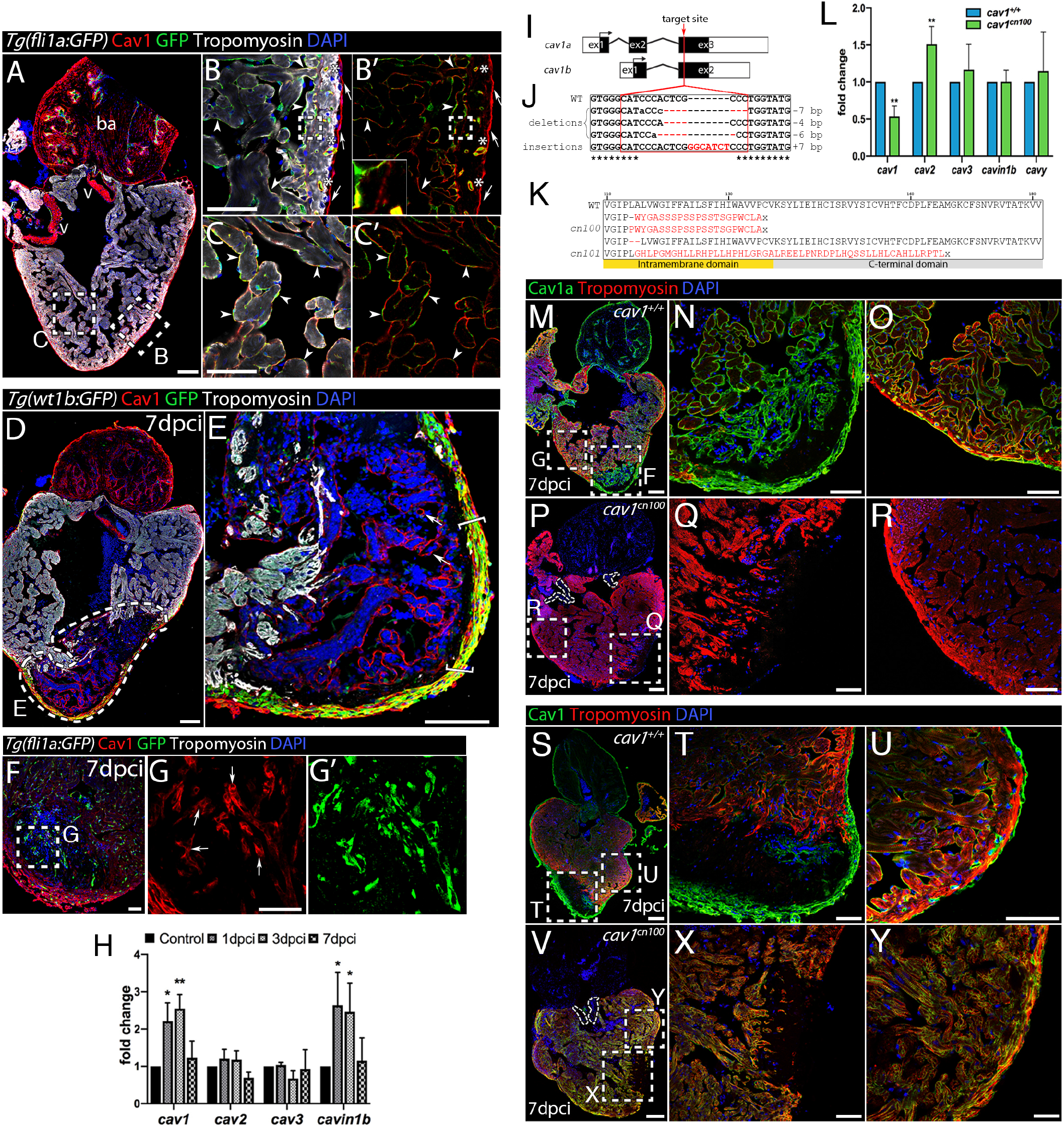
Caveolin-1 expression in the endothelium, endocardium and epicardium of the intact and injured adult zebrafish heart is abolished in CRISPR/Cas9 generated *cav1*-KO mutants. (A-C΄΄) Immunostaining of Cav1 and Tropomyosin (cardiomyocytes) in intact *Tg(fli1a:GFP)* heart. ba=bulbus arteriosus, v=valves. (B-C΄΄) Cav1 immunoreactivity in the epicardium (red, arrows) overlaps with GFP in the endothelium (asterisks) and endocardium (arrowheads). Cav1 is also expressed in the zone between the cortical and trabecular cardiomyocytes (dashed area in B-B΄, inset in B΄). (D,E) Cav1 immunostaining in 7dpci *Tg(wt1b:GFP)* heart. The dashed area denotes the injured area; Cav1 is expressed in the activated epicardium (E, brackets) and endocardium (E, arrows) upon injury. (F-G΄) Immunolabelling of Cav1 in 7dpci *Tg(fli1a:GFP)* heart. (G-G΄) Arrows indicate Cav1^+^ endocardial cells. (H) qPCR analysis of caveolae-related genes during regeneration. Mean±s.d., Brown-Forsythe and Welch ANOVA tests, \**P*<0.05, \*\**P*<0.01. Schematic representation of the two *cav1* transcripts, *cav1a* and *cav1b*. Arrowhead=target site, ex=exon. Identified mutations. Red dashed lines indicate deletions, red uppercase letters insertions, and black lowercase letters silent mutations. bp=base pairs. Cav1 domain organisation and the predicted effect on the protein. Red letters=novel amino acids, x=stop codon. qPCR analysis of caveolae-related genes in 2-day post fertilisation (dpf) embryos. Mean±s.d., t-test, \**P*<0.05, \*\**P*<0.01, \*\*\**P*<0.001. (M-R) Cav1a immunostaining in 7dpci *cav1*^*+/+*^ (M-O) and *cav1*^*cn100*^ (P-R) hearts. (S-Y) Cav1 (Cav1 and Cav1b) immunolabelling in 7dpci *cav1*^*+/+*^ (S-U) and *cav1*^*cn100*^ (V-Y) hearts. Dotted lines in P and V mark the valves. Scale bars 100μm in A, D, E, M, P, S, V and 50μm in other panels.

### Generation of *cav1*-KO zebrafish by CRISPR/Cas9 editing

We generated *cav1-KO* zebrafish to study the role of Cav1 and caveolae in heart homeostasis and regeneration. We used CRISPR/Cas9 editing to target the third exon of *cav1*, which corresponds to the C-terminal domain (Fig. 1I). The *cav1* gene generates two transcripts -*cav1a* and *cav1b*- sharing the majority of the coding sequences (Fang et al., 2006). We were able to introduce mutations in the *cav1* locus, with the majority of them being small deletions (Fig. 1J). The predicted effect on the protein was an open reading frame shift that would lead to an amino acid change and the generation of a premature stop codon (Fig. 1K). We selected the *cav1*^*cn100*^ mutation (Fig. 1K), which showed significantly decreased *cav1* expression, increased *cav2* expression, but with no effect on *cav3*, *cavin1b* or *cavy* transcription (Fig. 1L). Mutant embryos had no morphological abnormalities, developed normally and were fertile (data not shown). We next investigated Cav1 expression by labelling 7dpci WT and mutant hearts with an antibody against Cav1a (Fig. 1M-R). The strong Cav1 signal in the epicardium, endocardium, endothelium, bulbus arteriosus and valves (Fig. 1M-O) was lost in *cav1*^*cn100*^ hearts, indicating the loss-of-function nature of the mutation (Fig. 1P-R). We repeated this analysis with an antibody that recognises both Cav1 proteins, Cav1a and Cav1b (Fig. 1S-Y). Normal Cav1 expression in epicardium, endocardium and endothelium (Fig. 1S-U) was absent in mutants (Fig. 1V-Y). Cavin1 is also an essential component of caveolae (Hill et al., 2008) and deletion of Cav1 diminishes Cavin1 expression in mice (Hansen et al., 2013). We therefore asked whether Cavin1 was affected by Cav1 loss (Fig. S1). Examination of 7dpci *cav1*^*+/+*^ cryoinjured hearts revealed strong Cavin1 expression in the epicardium, in cardiomyocytes adjacent to the injured area, in the bulbus arteriosus and in the valves (Fig. S1A-D΄). Cavin1 expression was decreased in *cav1*^*cn100*^ hearts (Fig. S1E-H΄). Specifically, Cavin1 was absent in the valves, whereas its expression was greatly reduced in the bulbus arteriosus (Fig. S1E) and in cardiomyocytes within the proliferative zone (Fig. S1G-G΄). Taken together, these results show that the *cav1*^*cn100*^ mutation leads to the loss of Cav1 and the reduced expression of Cavin1 in cryoinjured hearts.

### Loss of caveolae in *cav1^cn100^* mutant hearts

Expression of *Cav1* in a system without endogenous Cav1 expression leads to *de novo* caveolae formation (Fra et al., 1995), whereas deletion of *Cav1* results in loss of caveolae (Drab et al., 2001). We analysed *cav1*^*+/+*^ and *cav1*^*cn100*^ hearts by transmission electron microscopy (TEM) to determine whether *cav1*^*cn100*^ mutants could form caveolae (Fig. S2). We found caveolae in abundance in the *cav1*^*+/+*^ hearts (Fig. S2A, A΄, C), and the membrane of the endothelial cells was packed with caveolae-invaginations (Fig. S2A΄, arrowheads). By contrast, *cav1*^*cn100*^ hearts were deprived of caveolae in the coronary vasculature of the cortical layer (Fig. S2B, B΄, C). No membrane-bound caveolae were detected, indicating the complete loss of caveolae in *cav1*^*cn100*^ mutant hearts.

### Response of caveolae-depleted hearts to injury

We next examined the effects of caveolae loss for adult heart regeneration. We cryoinjured *cav1*^*+/+*^ and *cav1*^*cn100*^ hearts and allowed them to regenerate for 90 days. We then analysed the hearts by Acid Fuchsin Orange-G (AFOG) staining, which labels both the damaged area and the healthy myocardium. We found that *cav1*^*cn100*^ and *cav1*^*cn100/+*^ hearts regenerated similarly to the *cav1*^*+/+*^ hearts 90dpci (Fig. 2G-J). Cryoinjury results in the formation of a scar tissue that progressively degrades during the course of 90 days (Chablais and Jaźwińska, 2012; González-Rosa et al., 2011; Schnabel et al., 2011). Thus, we monitored the regeneration process by analysing 30 and 60dpci hearts (Fig. 2A-F). *cav1*^*cn100*^ hearts had a similar scar size to that in *cav1*^*+/+*^ controls, both at 30 and 60dpci.

**Figure 2.**
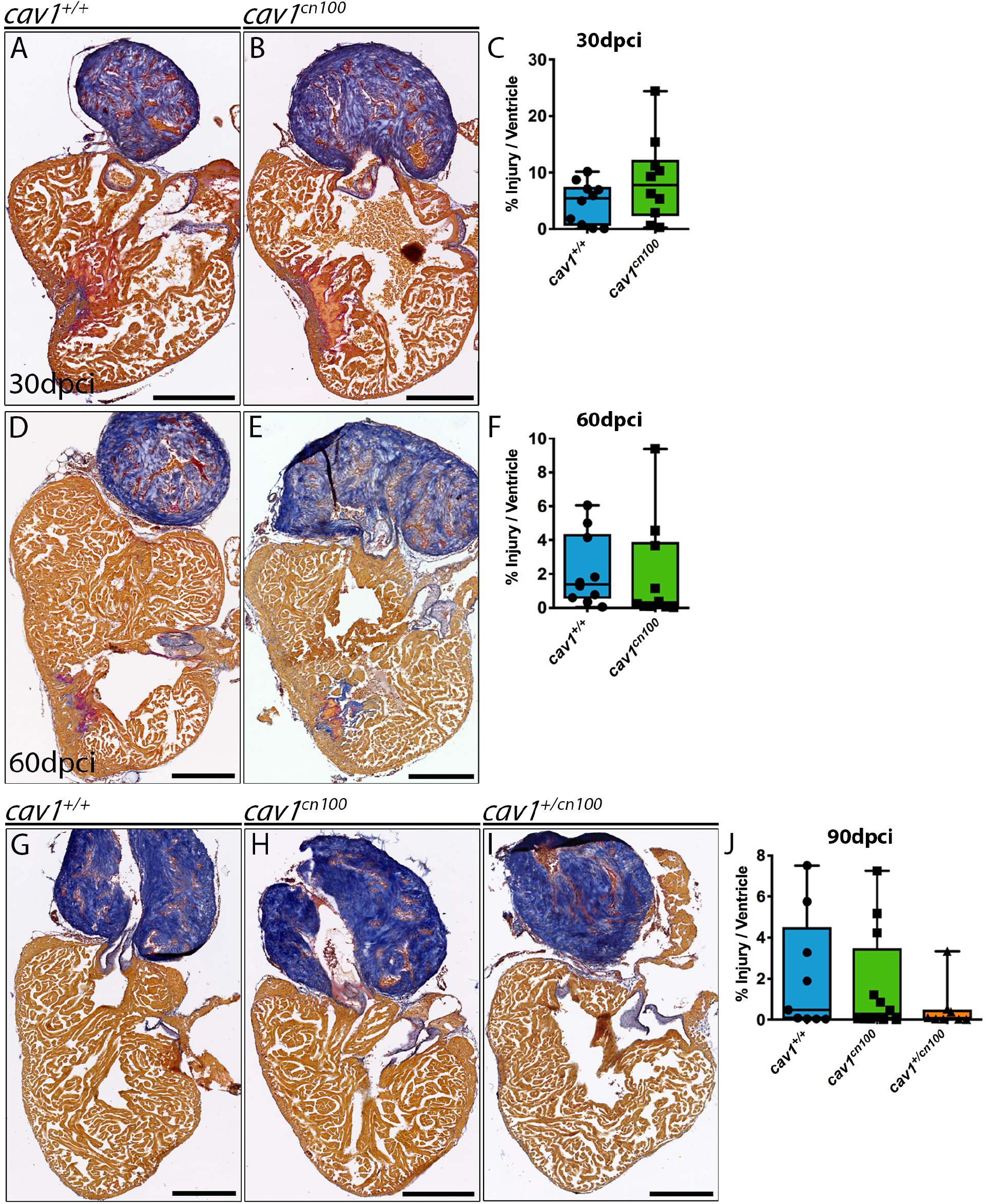
Heart regeneration is unaffected in *cav1*^*cn100*^ mutants. (A-J) *cav1*^*+/+*^ and *cav1*^*cn100*^ hearts were cryoinjured and harvested 30 (A-C), 60 (D-F), or 90dpci (G-J), and processed for AFOG staining, which labels collagen in blue, fibrin in red and myocardium in brown. Injuries were quantified as a percentage of the damaged tissue (collagen and fibrin) to the total area of the ventricle. 30dpci n_WT_=n_cn100_=10; 60dpci n_WT_=n_cn100_=10; 90dpci n_WT_=9, n_cn100_=12, t-test. Scale bars 250μm.

To further investigate the effect of caveolae loss in heart regeneration, we examined another of our *cav1* mutants, *cav1^cn101^*. Interestingly, *cav1* expression in *cav1*^*cn101*^ mutant embryos was unchanged, whereas *cav2* was upregulated (Fig. S3A). In addition, the *cav1*^*cn101*^ mutation had the same effect on protein expression as the *cav1*^*cn100*^ mutation, with loss of Cav1 expression (Fig. S3B-M). Likewise, caveolae were absent in *cav1*^*cn101*^ hearts, similar to our observations for *cav1*^*cn100*^ (Fig. S4). We then cryoinjured *cav1*^*+/+*^ and *cav1*^*cn101*^ hearts and examined the regeneration process every 30 days for 90 days (Fig. S5). *cav1*^*cn101*^ hearts regenerated normally and we did not detect any differences in the size of the injury at 30, 60 or 90dpci.

These results demonstrate that loss of caveolae does not affect heart regeneration. This was unexpected, as it has been reported that ventricular resection in *cav1-KO* zebrafish results in enhanced fibrosis, leading to the arrest of heart regeneration at 30dpci (Cao et al., 2016). The main difference between the ventricular resection and the cryoinjury models is that the fibrotic scar is minimal in the former, whereas the latter leads to formation of a transient scar tissue. Our data indicate that formation and resolution of the fibrotic tissue scar is unaffected by Cav1 loss. Full regeneration after ventricular resection occurs 60 days post amputation (Poss et al., 2002), and it would be worth to examine heart regeneration at the final time point. In addition, despite our *cav1* mutants harbour loss-of-function alleles, similar to those generated by Cao et al., 2016, the possibility of a genetic compensation mechanism (El-Brolosy et al., 2019) cannot be discarded.

### Activation of TGFβ signalling and fibrosis are unaffected in *cav1^cn100^* hearts

Caveolae are involved via Cav1 in the regulation of the TGFβ pathway (Razani et al., 2001), a major signalling node that controls extracellular matrix (ECM) deposition, and is activated during zebrafish heart regeneration (Chablais and Jazwinska, 2012). As both *cav1*^*cn100*^ and *cav1*^*cn101*^ hearts regenerated normally, we focused only on *cav1^cn100^*. To address TGFβ activity in *cav1*^*cn100*^ hearts upon cryoinjury, we quantified the nuclear localization of phospho-Smad3 (psmad3), a downstream effector of TGFβ (Fig. S6). We used 14dpci *Tg(fli1a:GFP)* hearts to calculate the proportion of psmad3^+^ nuclei in endocardial cells within the damaged tissue (Fig. S6A΄, B΄, C) and in the cardiomyocytes surrounding the injured site (Fig. S6A΄΄, B΄΄, D). Analysis revealed that TGFβ signalling was equally active in control and *cav1*^*cn100*^ hearts, in both endocardial cells and cardiomyocytes. Heart cryoinjury in the *Cav1-*KO mouse leads to extensive collagen deposition and cardiac remodelling (Miyasato et al., 2011). Thus, we exploited the AFOG staining protocol to examine the collagen and fibrin content after injury, and we also measured ventricular size (Fig. S7A, B). We found no differences between *cav1*^+/+^ and *cav1*^*cn100*^ hearts, neither in collagen deposition nor in fibrin amount in the injury, nor in the size of the ventricle. Furthermore, because hearts of *Cav1-KO* mice show increased interstitial fibrosis (Cohen et al., 2003; Drab et al., 2001; Murata et al., 2007; Park et al., 2003), we examined this parameter in intact *cav1*^*cn100*^ hearts using Picrosirius Red to stain collagen fibres. Results showed no difference in interstitial fibrosis between *cav1*^*+/+*^ and *cav1*^*cn100*^ hearts (Fig. S7C-E). Thus, loss of Cav1 and caveolae does not affect TGFβ activity or fibrosis in intact or cryoinjured hearts.

### Epicardial, endocardial and cardiomyocyte function in *cav1^cn100^* hearts upon injury

We next examined the behaviour of the different cell types involved in the regeneration process. We first analysed the epicardium and endocardium, where Cav1 is highly expressed 7dpci (Fig. S8). We crossed the *Tg(wt1b:GFP)* line to *cav1*^*cn100*^ to study epicardial proliferation and we found no difference in epicardial proliferation between *cav1*^*+/+*^ and *cav*^*cn100*^ hearts (Fig. S8A-C).

We then evaluated the abundance of endocardial cells within the damaged tissue by crossing double transgenic fish *Tg(fli1a:GFP)/Tg(myl7:mRFP)* expressing GFP in endocardial/endothelial cells and RFP in the membrane of cardiomyocytes, with *cav1*^*cn100*^ mutants (Fig. S8D,E). Three-dimensional volume rendering and analysis of the GFP^+^ cells inside the RFP^-^ area showed that endocardial cells in *cav1*^*cn100*^ hearts populated the injured area similarly to those of *cav1^+/+^*hearts (Fig. S8F). These data indicate that epicardial proliferation and migration and proliferation of endocardial cells are unchanged in *cav1*^*cn100*^ hearts.

We then examined cardiomyocyte proliferation, as it has been reported that this feature is affected upon ventricular resection in a *cav1*-KO zebrafish model (Cao et al., 2016). We analysed the cardiomyocytes adjacent to the injured area 7dpci, because they are highly proliferative (Bednarek et al., 2015). Quantification of BrdU incorporation into cardiomyocytes revealed that proliferation was significantly lower in *cav1*^*cn100*^ hearts than in *cav1*^*+/+*^ hearts (Fig. 3A-C). We confirmed this result in the *cav1*^*cn101*^ animals (Fig. S9). We then addressed the proliferation status of cardiomyocytes at 14dpci (Fig. 3D-F). We found that at this time point, *cav1*^*cn100*^ cardiomyocytes proliferated at the same rate as *cav1*^*+/+*^ cardiomyocytes. These data indicate that loss of caveolae leads to the transient attenuation in *cav1*^*cn100*^ cardiomyocyte proliferation at 7dpci, which is normalised by 14dpci, leading to normal cardiac regeneration at 90dpci.

**Figure 3.**
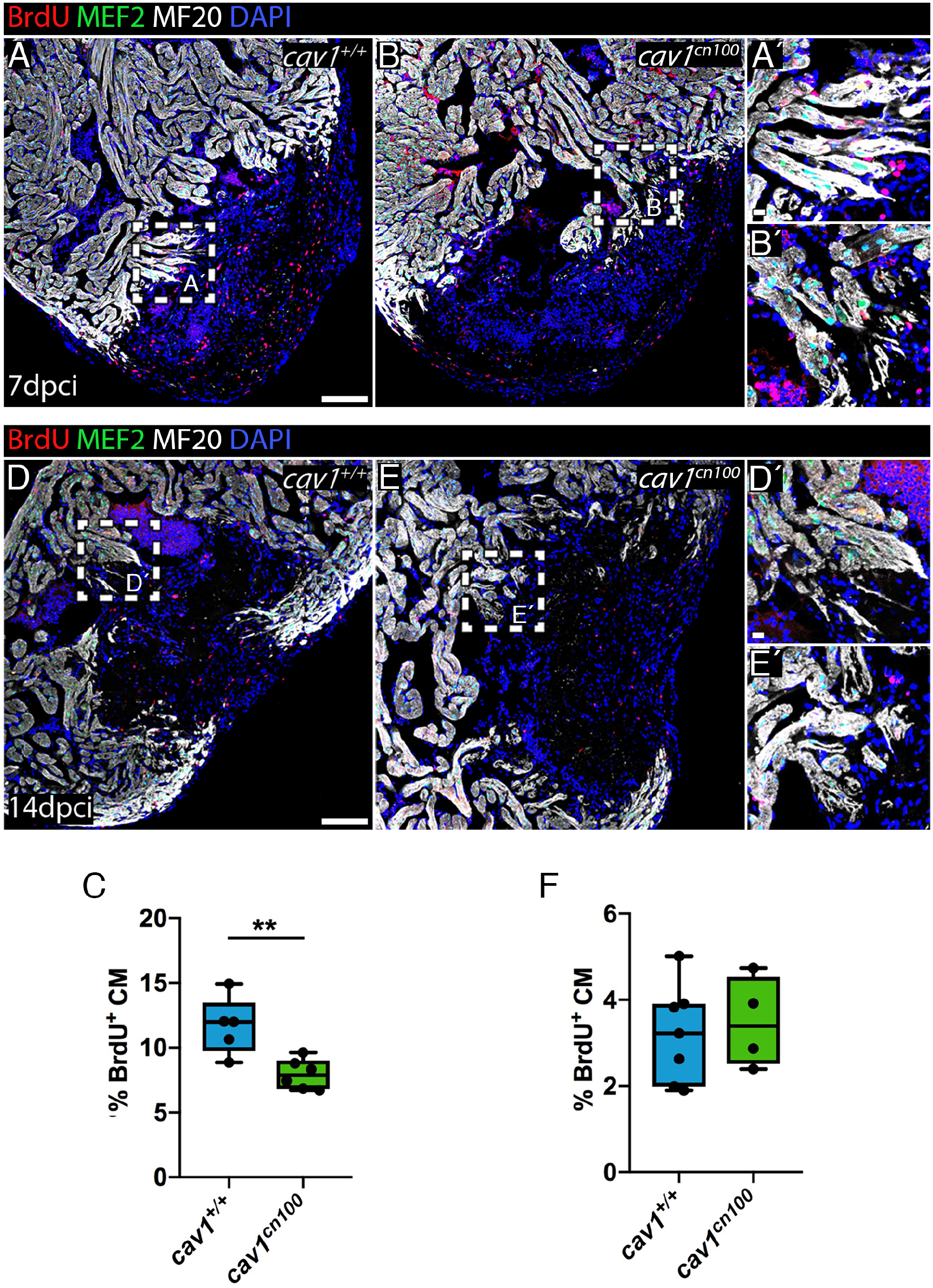
Cardiomyocyte proliferation is transiently reduced upon cryoinjury in *cav1*^*cn100*^ hearts. (A-B) Immunolabelling of 7dpci *cav1*^*+/+*^ and *cav1*^*cn100*^ hearts for BrdU, MEF2 (cardiomyocyte nuclei) and MF20 (cardiomyocytes). (A΄-B΄) magnifications of the dashed areas in A and B. (C) Quantification of the BrdU^+^/MEF2^+^ nuclei to the total cardiomyocyte number in a 100μm radius around the injured area. n_WT_=5, n_cn100_=6, t-test, \*\**P*<0.01. (D-E) Immunolabelling of 14dpci *cav1*^*+/+*^ and *cav1*^*cn100*^ hearts for BrdU, MEF2 and MF20. (F) Quantification of cardiomyocyte proliferation at 14dpci. n_WT_=7, n_cn100_=4. Scale bars: 100μm in A, B, D, E; 10μm in A΄, B΄, D΄, E΄.

### Impaired heart function and cardiac elasticity in *cav1^cn100^* mutants

*Cav1*-deficient mice show decreased systolic function (Cohen et al., 2003; Zhao et al., 2002). Therefore, we analysed heart function in our mutants using ultrasound imaging. Echocardiography of adult *cav1*^*+/+*^ and *cav1*^*cn100*^ animals revealed that *cav1*^*cn100*^ hearts were less efficient in pumping blood than control hearts, as their ejection fraction was significantly lower (Fig. 4A). Additionally, the heart rate of *cav1*^*cn100*^ animals was lower than in control siblings (Fig. 4B). These findings indicate that caveolae are essential for normal cardiac function.

**Figure 4.**
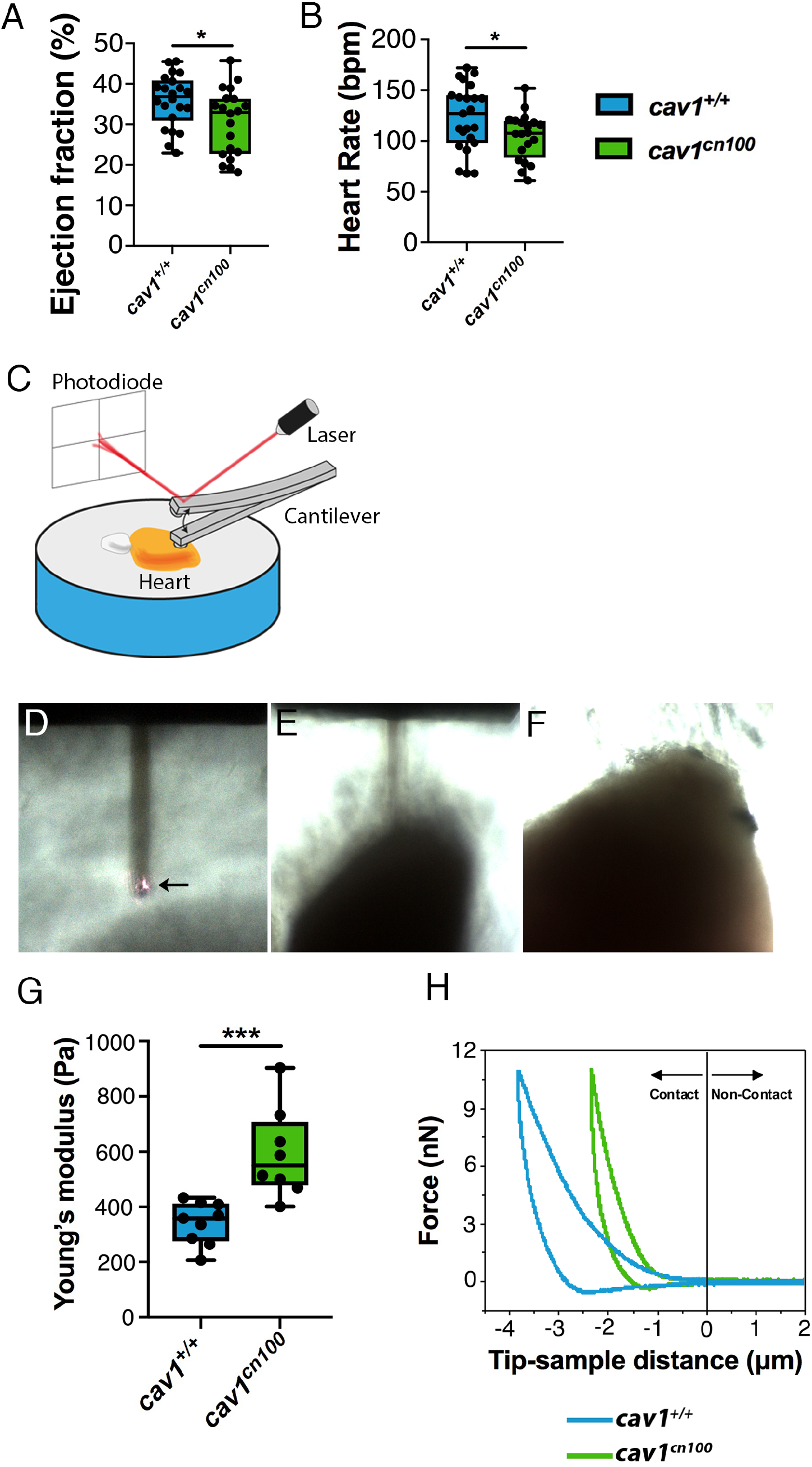
Impaired cardiac function and stiffer heart tissue in caveolae-deprived *cav1*^*cn100*^ mutants. (A) Quantification of ejection fraction in *cav1^+/+^* and *cav1^cn100^* hearts. n_WT_=22, n_cn100_=20. (B) Heart rate measurements in *cav1^+/+^* and *cav1^cn100^* animals. n_WT_=23, n_cn100_=20. t-test, \**P*<0.05. (C) AFM set-up. Arrow indicates the cantilever and the laser beam. (D-F) The ventricular apex was oriented towards the cantilever and used for taking measurements. (G) Biomechanical characterization of *cav1*^*+/+*^ and *cav1*^*cn100*^ ventricles as measured by AFM-force spectroscopy and expressed in Young’s moduli. n_WT_=9, n_cn100_=8, t-test, \*\*\**P*<0.001. (H) Force-distance graph for indentation and retraction of the cantilever over the ventricular surface.

Caveolae provide protection against mechanical stress (Sinha et al., 2011). Because the biomechanical properties of cells influence the behaviour of tissues (Mathur et al., 2001) and caveolae are involved in mechanoprotection, we investigated the mechanical response of *cav1*^*+/+*^ and *cav1*^*cn100*^ cardiac tissue to gain insight into the differences observed in cardiac function. We used AFM to determine the response of adult *cav1*^*cn100*^ hearts to force. Freshly isolated hearts were placed on an agarose gel, with the apex of the ventricle oriented towards the direction of the cantilever, and force measurements were taken (Figure 4C-F). We found that cardiac tissue stiffness was significantly greater in *cav1*^*cn100*^ animals than in control animals (Figure 4G). Applying the same force, the extent of deformation was 1.5 times greater in control hearts than in *cav1*^*cn100*^ hearts, without causing permanent deformation as the curves returned to zero (Figure 4H).

Importantly, the sensitivity of the method revealed that changes in epicardial and cortical myocytes were responsible for the observed difference in stiffness that we detected. Membrane invaginations such as caveolae, provide the necessary stretch capacity for cells to buffer the impact of mechanical forces (Sinha et al., 2011). Hence, the absence of membrane reservoirs, caveolae, due to the loss of the broad and robust expression of Cav1 in the ventricle, results in increased tissue stiffness. As we did not detect cardiac fibrosis, the reduction in cardiac elasticity is a plausible explanation for the cardiac dysfunction in *cav1-KO* fish.

We have shown that Cav1 is strongly expressed in the epicardium of intact hearts and that its expression is lost in *cav1*^*cn100*^ hearts. Additionally, RNA-seq data show that Cav1 expression is moderate but significantly higher in cortical cardiomyocytes, but significantly higher than in trabecular cardiomyocytes (Sánchez-Iranzo et al., 2018). Caveolae are required for smooth muscle cell contractility (Gosens et al., 2011; Halayko et al., 2008) and cell contraction results in changes to membrane tension that caveolae buffer (Sinha et al., 2011). In addition, Cav1 and caveolae regulate RhoA activation (a GTPase protein that regulates the cytoskeleton), which in turn regulates actomyosin contractility (Budnar et al., 2019; Grande-García et al., 2007; Peng et al., 2007). Accordingly, these data suggest that loss of Cav1 and caveolae impairs cell elasticity and/or contractility and, consequently, pumping efficiency, which could explain the significant reduction in cardiac performance. Cav1 deficiency in the mouse leads to diverse cardiac phenotypes that are attributed to endothelial loss of caveolae. Our results demonstrate that loss of Cav1 and caveolae result in cardiac stiffening accompanied by reduced cardiac function, suggesting that a global change in the mechanical properties of the heart leads to the cardiac phenotypes observed in Cav1-deficiency models.

## Material and Methods

### Zebrafish husbandry and transgenic lines

Animal studies were approved by the CNIC Animal Experimentation Ethics Committee and by the Community of Madrid (Ref. PROEX 118/15). Animal procedures conformed to EU Directive 2010/63EU and Recommendation 2007/526/EC regarding the protection of animals used for experimental and scientific purposes, enforced in Spanish law under Real Decreto 53/2013. Zebrafish were raised under standard conditions at 28ºC as described (Kimmel et al., 1995). Experiments were performed with 5-14-month-old adults.

### CRISPR/Cas9 injections

We used the oligos *cav1* Fwd CACCGGTGGGCATCCCACTCGCCC and *cav1* Rvs AAACGGGCGAGTGGGATGCCCACC for the generation of *cav1-KO* zebrafish. The oligos were inserted into the pX330-U6-Chimeric_BB-CBh-hSpCas9 vector (Ann, F et al., 2013; Cong et al., 2013), which was linearised with BbsI enzyme (New England Biolabs, Ipswich, MA). Primers with the T7 polymerase promoter-specific sequences, *cav1* T7 Fwd TAATACGACTCACTATAGGTGGGCATCCCA and Rvs AAAAAGCACCGACTCGGTGCCA, were used to amplify the guide RNA, which was injected into one-cell state embryos together with Cas9 protein (New England Biolabs). Mutant animals were identified by PCR using the primers: *cav1* Fwd GGCGAGCTTCACCACCTTC and *cav1* Rvs GCTCTTCACGCAAGGCACCA.

### Adult heart cryoinjury

Fish were anesthetised by immersion in 0.04% tricaine (Sigma-Aldrich, St Louis, MO) in fish water and placed on a wet sponge under a stereoscope with the ventral side exposed. The cardiac cavity was opened using microscissors and microforceps, and the pericardium was removed. The ventricle of the heart was exposed and dried and was then touched by a copper-made probe previously immersed in liquid nitrogen (González-Rosa and Mercader, 2012). The fish were immediately returned to water to recover.

### Bromodeoxyuridine injection

Adult fish were anesthetised and placed on a wet sponge under a stereoscope. BrdU was diluted in phosphate-buffered saline (PBS) to 2.5 mg/ml and 30 μl were injected intraperitoneally.

### Echocardiography

Analysis of cardiac function by echocardiography in adult fish was performed as described (González-Rosa et al., 2014). Briefly, the fish were anesthetised by immersion in 60 mM tricaine and 3 mM isoflurane in fish water and transferred to a sponge immersed in the same solution. Images were acquired using the VEVO 2100 system (VisualSonics Inc., Toronto, ON, Canada) with a 50-MHz ultrasound probe. The transductor was immersed in the medium dorsally to the cardiac cavity. The fish were immediately transferred to fresh water to recover after the procedure.

### Histological stains

Acid Fuchsin Orange G-staining (AFOG) and Picrosirius Red staining were performed following standard protocols (Poss et al., 2002).

### Immunofluorescence

Sections of paraffin-embedded tissue were permeabilised with PBT (PBS with 0.01% TritonX-100) and washed with PBS before incubation with blocking solution (2% bovine serum albumin, 10% goat serum and 2 mM MgCl_2_ in PBS). Sections were then incubated overnight at 4ºC with the following primary antibodies: Caveolin-1 (BD Transduction Laboratories, San Jose, CA; or Cell Signalling Technology, Danvers MA), PTRF/Cavin1 (Atlas Antibodies AB, Stockholm, Sweden), GFP (Aves Labs, Tigard, OR), MEF-2 (Santa Cruz Biotechnology, Santa Cruz, CA), tropomyosin, MF20 (both from DSHB, Iowa City, IA), phosho-Smad3 (Abcam, Cambridge, MA) and BrdU (BD Transduction Laboratories) The following day, sections were incubated with the appropriate secondary antibody and mounted after DAPI staining.

### Whole-mount confocal imaging

Analysis of endogenous fluorescence of whole-mount hearts was performed as described (Münch et al., 2017). Briefly, hearts were fixed overnight with 2% paraformaldehyde and after several PBS washes the tissues were immersed in 3% agarose. Samples were then incubated in CUBIC I solution (Susaki et al., 2014) at 37ºC for one week. Agarose blocks containing the hearts were mounted for imaging on a petri dish and approximately 700μm of the injured ventricle was scanned on a Leica SP8 confocal microscope (Leica Microsystems, Wetzler, Germany) using a 10× objective.

### Electron microscopy

Hearts were fixed in 1% glutaraldehyde/4% paraformaldehyde in PBS overnight. Samples were post-fixed in 1% osmium tetroxide for 60 min and dehydrated through a series of ethanol solutions (30%, 50%, 70%, 95%, 100%) and acetone. After the last dehydration step, samples were incubated in a 1:3, 1:1, 3:1 mixture of DURCUPAN resin and acetone and cured at 60ºC for 48 hours. Ultrathin sections (50-60nm) were obtained using a diamond knife (Diatome AG, Biel, Switzerland) in a ultramicrotome (Leica Reichert ultracut S. Leica Microsystems) and collected in 200-mesh copper grids. The sections were counterstained with 2% uranyl acetate in water for 20 min followed by a lead citrate solution. Sections were examined with a JEOL JEM1010 electron microscope (Tokyo, Japan) equipped with an Orius SC200 digital camera (Gatan Inc., Pleasanton, CA).

### Image analysis and quantification

To analyse cardiomyocyte proliferation, MEF2-positive nuclei were counted in an area of 100μm around the injury site using Fiji (ImageJ, NIH). BrdU-MEF2-positive cells were also counted and the % proliferation index was expressed as the MEF2:BrdU-MEF2 ratio. For TGFβ signalling activation, all phospho-smad3-positive cardiomyocyte (100μm of the injury) or GFP-positive cells (inside the injured area) were counted and normalised to the total number of cardiomyocytes or GFP-positive cells. To quantify the regeneration process, the injured area (fibrotic tissue and collagen) was measured using Fiji and expressed as a percentage of the total ventricular area. For tissue sections, at least three sections per sample were used for measurements. The 3D analysis of the whole-mount hearts was carried out using Fiji and IMARIS programmes. The volume of the GFP signal inside the injured area (RFP negative) was also measured and presented in relation to the volume of the injury. Electron microscopy images of the plasma membrane of coronary endothelial cells were taken at 50,000× magnification. Uncoated membrane invaginations of 40-90nm size were counted (Lim et al., 2017) and expressed as density per μm^2^ of the perinuclear area. Two endothelial cells per three sections of the same heart were examined. Fiji was used to calculate the perinuclear area and for caveolae identification.

### Atomic force microscopy (AFM)-force spectroscopy

Adult zebrafish were sacrificed by immersion in 0.16% tricaine and the heart was dissected. The atrium was removed and the ventricle was placed horizontally atop a 4% agarose gel immersed in PBS with 0.1M KCl to arrest the heartbeat uniformly. AFM-force spectroscopy experiments were performed with a JPK Nanowizard III microscope (JPK Instruments, Berlin, Germany) coupled with an inverted optical microscope (AXIO Observer D1; Carl Zeiss, Germany) and equipped with Plateau-CONT-SPL cantilevers (Nanosensors, Neuchatel, Switzerland), with a nominal spring constant of 0.02-0.77 N/m and a spherical tip shape (R = 30 μm). The actual spring constant of the cantilever was determined using the thermal noise method as implemented in the AFM software. Force-distance curves (FDC) were acquired to determine Young’s modulus of the zebrafish heart apex. The tip-sample distance was modulated by applying a triangular waveform (Garcia et al., 2017b). The tip velocity was set to 10 μm/s and the amplitude to 15μm. The maximum force exerted on the heart apex during a single FDC was of 11 nN. For each zebrafish heart, a complete sequence of 375 FDC was performed. These FDC were distributed in three different areas of 100×100 μm^2^ several hundreds of microns apart. In each area, 125 FDC were measured (all along the zebrafish apex). To determine the contact point, we used a ratio of variances protocol. Young’s modulus was obtained by fitting a section of the force-distance curve (approach semi-cycle of the whole FDC) with a Hertz model for spherical indenters.

### Quantitative RT-PCR

Three-to-five biological replicates with three technical replicates of each sample were used for the expression analysis of genes by qPCR using the power SYBR Green Master Mix (Applied Biosystems, Foster City, CA) and the ABI PRISM 7900HT FAST Real-Time PCR System. All measurements were normalised to the expression of 18s (McCurley and Callard, 2008). The following primers were used for the qPCR analysis: 18s Fwd TCGCTAGTTGGCATCGTTTATG, 18s Rvs CGGAGGTTCGAAGACGATCA, cav1 Fwd TGGGATGGGGGAATGGAAAC, cav1 Rvs TAAACGGCGAGTGAGCGTAT, cav2 Fwd GCGTTTATTGCAGGGATTGT, cav2 Rvs GGATCACTGGCATCACCAC, cav3 Fwd CAACGAAGATGTCGTGAAGG, cav3 Rvs GAGACGGTGAAGGTGGTGTAA, and for cavin1b and cavy form (Lim et al., 2017).

### Statistical analysis

Sample sizes, statistical tests and P-values are specified in the figure legends and were determined with GraphPad Prism software (GraphPad Software Inc., San Diego, CA).

## ACKNOWLEDGMENTS

We thank E. Dı́az at the CNIC animal facility for fish husbandry; B. Rios, V. García, L. Méndez for technical support; the CNIC Microscopy Unit for help; A.Vanesa Alonso and L. Flores for support with the echocardiographic experiment; F. Urbano and C. Aguado for support and help with the TEM.

---

**Figure S1. Cavin1b expression in***cav1*^*cn100*^ **hearts**

Immunostaining of Cavin1 in 7dpci *cav1*^*+/+*^ (A-D΄) and *cav1*^*cn100*^ (E-H΄) *Tg(wt1b:GFP)* hearts. (B) Magnification of selected area in A. (C-D΄) Magnifications of the dashed areas in B. (F) Magnification of selected area in E. (G-H΄) Magnifications of the dashed areas in F. (A, E) Dashed lines=valves. Scale bars 100μm in A, B, E, F; 50μm in other panels.

**Figure S2. Loss of caveolae in***cav1*^*cn100*^ **hearts**

TEM images of *cav1*^*+/+*^ (A, A΄) and *cav1*^*cn100*^ (B, B΄) endothelium. (A΄, B΄) Higher magnification of the dashed areas in A and B. Arrowheads indicate membrane-bound caveolae. Scale bars: 1μm in A, B; 0.5μm in A΄, B΄.

(C) Caveolae number per μm^2^ of endothelial cell. n_WT_=n_cn100_=4, t-test, \*\**P*<0.01.

**Figure S3. qPCR and Cav1 expression analysis in***cav1*^*cn101*^ **mutants**

(A)Relative expression of caveolae-related genes by qPCR in *cav1*^*cn101*^ embryos. Mean±s.d. t-test, \*\*\**P*<0.001.

(B-M) Cav1a (B-G) or Cav1 (H-M) immunostaining of 7dpci *cav1*^*+/+*^ and *cav1*^*cn101*^ hearts. Scale bars 100μm B, E, H, K and 50μm in other panels.

**Figure S4. Caveolae loss in***cav1*^*cn101*^ **hearts**

(A-B΄) Coronary vasculature of the cortical layer in *cav1*^*+/+*^ and *cav1*^*cn101*^ hearts. A΄ and B΄ higher magnifications of the dashed areas in A and B; arrowheads indicate membrane-bound caveolae. Scale bars: 1μm in A, B, and 0.5μm in A΄ and B΄.

Quantification of caveolae number per μm^2^ of coronary endothelium. n_WT_=n_cn100_=4, Mean±s.d., t-test, \*\*\*\**P*<0.0001.

**Figure S5. Heart regeneration is unaffected in***cav1*^*cn101*^ **mutants**

(A-J) AFOG staining and quantification in *cav1*^*+/+*^ and *cav1*^*cn101*^ hearts after 30 (A-C), 60 (D-F) and 90dpci (G-J). Collagen in blue, fibrin in red and myocardium in brown. The damaged area was quantified as the percentage of the collagen/fibrin area to the total ventricular area. 30dpci n_WT_=9, n_cn100_=7; 60dpci n_WT_=8, n_cn100_=7; 90dpci n_WT_=7, n_cn100_=11. Mean±s.d., t-test. Scale bars: 200μm.

**Figure S6. TGFβ signalling activation is unaffected in regenerating***cav1*^*cn100*^ **hearts**

(A-B) 14dpci *cav1*^*+/+*^ and *cav1^cn100^ Tg(fli1a:GFP)* hearts labelled for psmad3, GFP and MF20 (cardiomyocytes). (A΄, B΄) Magnification of GFP^+^ endocardial cells marked in A, B. (A΄΄, B΄΄) Higher magnifications of cardiomyocytes marked in A, B. (C) Quantification of the psmad3^+^/GFP^+^ in the injured area. t-test. n_WT_=7, n_cn100_=4. (D) Percentage of cardiomyocytes with psmad3^+^ nuclei in a 100μm area surrounding the damaged tissue. t-test, n_WT_=7, n_cn100_=4. Scale bars: 100μm A, B; 25μm in other panels.

**Figure S7. Analysis of collagen and fibrin of the injured area, ventricular size and interstitial fibrosis of***cav1*^*cn100*^ **hearts**

(A)Percentages of the collagen and fibrin of the injury 30-, 60- and 90dpci in *cav1*^*+/+*^ and *cav1^cn100^* hearts. Two-way ANOVA. n_WT_ 30-, 60-, 90dpci = 10, 9, 9; n_cn100_ 30-, 60-, 90dpci = 10, 10, 12.

(B)Ventricular size of all hearts analysed by AFOG staining. t-test. n_WT_=29; n_cn100_=32.

(C-D) Picrosirius Red staining in control *cav1*^*+/+*^ and *cav1*^*cn100*^ hearts. Scale bar 250μm.

(E) Quantification of the red-labelled fibres in the ventricle. t-test, n_WT_=5, n_cn100_=6.

**Figure S8. Normal epicardial proliferation and endocardial cell function in***cav1*^*cn100*^ **hearts after injury**

(A-B) Immunostaining of 7dpci *Tg(wt1b:GFP)* heart sections labelled for BrdU and GFP.

(C) Percentage of proliferating epicardial GFP^+^ cells. n_WT_=5, n_cn100_=6. Scale bar 100μm.

(D-E) 3D volume rendering of the apical injured site of 7dpci *Tg(fli1a:GFP)/Tg(myl7:mRFP)* hearts. Yellow lines indicate the injured area and heart cartoon the x/y/z axes.

(F) Quantification of the volume of GFP^+^ cells inside the RFP^-^ area. t-test, n_WT_=3, n_cn100_=4. Scale bar 300μm.

**Figure S9.***cav1*^*cn101*^ **cardiomyocyte proliferation upon injury**

(A-B΄) Immunolabelling of 7dpci *cav1*^*+/+*^ and *cav1*^*cn101*^ hearts for BrdU, MEF2 and MF20. (A΄-B΄) Higher magnifications of the dashed areas in A and B. Scale bars: 100μm in A, B; 20μm in A΄, B΄.

(C) Percentage of the BrdU^+^ cardiomyocytes to the total number of cardiomyocytes in a 100μm area surrounding the damaged tissue. n_WT_=4, n_cn100_=3. t-test, \**P*<0.05.

